# Characterization of growth and development of sorghum genotypes with differential susceptibility to *Striga hermonthica*

**DOI:** 10.1101/2021.02.24.432663

**Authors:** Dorota Kawa, Tamera Taylor, Benjamin Thiombiano, Zayan Musa, Hannah E. Vahldick, Aimee Walmsley, Alexander Bucksch, Harro Bouwmeester, Siobhan M. Brady

## Abstract

Variation in strigolactone composition in sorghum root exudates underlies its resistance to the parasitic weed, *Striga hermonthica*. Root exudates of the Striga susceptible variety Shanqui Red (SQR) contain primarily 5-deoxystrigol, which has a high efficiency of inducing Striga germination. SRN39 roots primarily exude orobanchol, leading to reduced Striga germination and making this variety resistant to Striga. This structural diversity in exuded strigolactones is determined by the polymorphism in the *LGS1* (*LOW GERMINATION STIMULANT 1*) locus. Yet, the effects of the *lgs1* mutation as well as the consequences of the vast genetic diversity between SQR and SRN39 have not been addressed in terms of growth and development. Here, we demonstrate additional consequences of *LGS1* loss-of-function by phenotypic and molecular characterization. A suite of genes related to metabolism was differentially expressed between SQR and SRN39. Increased levels of gibberellin precursors in SRN39 were accompanied with its slower growth rate and developmental delay and we observed an overall increased SRN39 biomass. The slow-down in growth and differences in transcriptome profiles of SRN39 were strongly associated with plant age. Additionally, analyses of multiple *LGS1* loss-of-function genotypes indicated that strigolactone stereochemistry influences root system architecture. In summary, we demonstrate that the consequences of the *lgs1* mutation reach further than the changes in strigolactone profile in the root exudate and translate into alterations in growth and development.

**Highlight:** SRN39 and Shanqui Red are models for sorghum genotypes that are resistant and susceptible, respectively, to *Striga hermonthica*. Additional differences in plant growth, development, and hormone abundance should be considered when assessing Striga tolerance.

## Introduction

*Sorghum bicolor* belongs to the five most important cereal crops, globally (Paterson *et al*., 2009). Its adaptations to drought, heat and low nutrient availability, made sorghum especially important for agriculture in sub-Saharan Africa, where it is a staple food, feed and forage (Mace *et al*., 2013). Among sorghum strategies to withstand abiotic stresses, establishing interactions with arbuscular mycorrhiza (AM) fungi improves phosphate acquisition (Neumann and George, 2004). To recruit AM fungi, plants secrete strigolactones into the soil, carotenoid-derived compounds that induce AM hyphal branching (Akiyama *et al*., 2005). Low nutrient concentration, primarily phosphate and nitrogen, induces strigolactone exudation (Jamil *et al*., 2013), which ensures AM fungi recruitment (Bouwmeester *et al*., 2007; Yoneyama *et al*., 2007). While this plant-fungal communication system is beneficial for sorghum performance in nutrient-depleted soils, the strigolactone signal can be hijacked by seeds of the parasitic “witchweed”, *Striga hermonthica* (Bouwmeester *et al*., 2007). Striga is an obligate parasite and strigolactones are essential for its seed germination (Bouwmeester *et al*., 2020; Spallek *et al*., 2013). Co-option of strigolactone signaling by Striga ensures it germinates only in the presence of a host plant. The ability to sense the presence of strigolactones allows Striga seeds to stay dormant for years and makes Striga an extraordinarily successful parasite. After strigolactone perception, germinated Striga forms a haustorium which penetrates the roots of a host plant and establishes a connection between its own xylem and the xylem of the host plant, to deprive the host of water and nutrients (Spallek *et al*., 2013). Striga infestation has a detrimental effect on a host plant and around 20 percent of sorghum yield is lost annually (Ejeta and Gressel, 2007).

Since strigolactones are essential for Striga parasitism, understanding their biosynthesis and functions is essential to combat Striga parasitism. More than 35 different strigolactones have been discovered across plant species, yet not all have the potential to induce Striga germination to the same extent (Wang and Bouwmeester, 2018). Among the four main strigolactones produced by sorghum, 5-deoxystrigol, sorgomol, sorgolactone and strigol have high Striga germination stimulant activity, while orobanchol is a low germination stimulant (Awad *et al*., 2006; Gobena *et al*., 2017; Mohemed *et al*., 2018). Two sorghum varieties, Shanqui Red (SQR) and SRN39, have been studied extensively in terms of the relationship between strigolactones and Striga resistance. SRN39 is a Striga resistant, improved sorghum variety released in Sudan, while SQR is a Chinese sorghum variety with high susceptibility to Striga (Satish *et al*., 2012). SQR root exudate contains 5-deoxystrigol as the dominant strigolactone and has a high efficiency of Striga germination induction (Gobena *et al*., 2017). SRN39 roots exude mainly orobanchol, leading to low Striga germination. This change in strigolactone exudation profile was associated with a deletion in the *LGS1* (*LOW GERMINATION STIMULANT 1*) locus (Gobena *et al*., 2017; Satish *et al*., 2012). LGS1 encodes a putative sulfotransferase, but its role in strigolactone biosynthesis is not yet understood.

Several other sorghum genotypes possess the *lgs1* mutation and consequently have low Striga germination-inducing activity (Bellis *et al*., 2020; Mohemed *et al*., 2018). However, the *LGS1* loss-of-function alleles are not prevalent in Striga prone regions and are absent from non-infested areas, which suggests potential trade-offs of *lgs1* mutations (Bellis *et al*., 2020). Decreased expression of photosynthesis-related genes in root and shoot tissue of *LGS1* loss-of-function genotypes is one of the potential costs of Striga resistance (Bellis *et al*., 2020). Moreover, elevated expression of genes involved in strigolactone biosynthesis in SRN39 roots indicates additional alterations of strigolactone profiles within this genotype, that were not previously captured in root exudate profiles (Bellis *et al*., 2020). To date, it is not known whether the strigolactone composition in the root and shoot tissue reflects that of the root exudate.

Strigolactones also act as an endogenous plant hormone and as such regulate plant root and shoot architecture (Aquino *et al*., 2020). Strigolactones repress shoot branching (Gomez-Roldan *et al*., 2008; Umehara *et al*., 2008) and promote secondary growth of the stem (Agusti *et al*., 2011). The potential of strigolactones to shape root system architecture depends on the availability of phosphate. Under phosphate-sufficient conditions, strigolactones inhibit lateral root formation while promoting lateral root primordia formation in a phosphate-limited environment (Ruyter-Spira *et al*., 2011). Next to their impact on organ growth and development, strigolactones also affect plant metabolism. Exogenous application of strigolactones induces biosynthesis of organic acids involved in mobilization of inorganic phosphate (Gamir *et al*., 2020). It remains unknown whether the influence of strigolactones on plant development and metabolism varies with respect to different strigolactone structural variants.

Given the pleiotropic effect of strigolactones, the influence of LGS1 on strigolactone profiles and photosynthetic efficiency, one could expect that the *lgs1* genotypes would show differences in growth and plant architecture. However, only a minor reduction in leaf area was observed in the *lgs1* mutant (Bellis *et al*., 2020), and the consequences of *LGS1* loss-of-function on sorghum growth and development were never characterized in detail. Here, we use transcriptome profiling to identify biological processes that differentiate SQR and the *lgs1* genotype, SRN39. We show that genes involved in metabolism have altered expression in SRN39 as compared to SQR and further explore the phenotypic variation between these two varieties. Our results suggest that the consequences of the genetic differences between SQR and SRN39 for growth rate depend on plant developmental stage. We associate the reduced SRN39 shoot growth relative to SQR with increased accumulation of gibberellin precursors. We further characterize the root system architecture of several l*gs1* genotypes and speculate that various strigolactone structural variants affect root system architecture. Our results suggest that the differences between SQR and SRN39 go beyond distinct susceptibility to Striga, and include differences in growth, development and root system architecture.

## Materials and Methods

### Plant material

Seeds of *Sorghum bicolor* var. Shanqui Red (SQR) were obtained from GRIN (https://www.ars-grin.gov). Birhan, Framida, Gobiye and SRN39 seeds were kindly donated by the Ethiopian Institute of Agricultural Research. For each experiment seeds were surface sterilized by agitating in a solution containing 30% (v/v) commercial bleach and 0.2% Tween-20 (v/v) for 30 minutes followed by five washes with sterile water and overnight incubation in 5% (w/v) Captan fungicide. Sterilized seeds were germinated on wet Whatman paper (grade 1) upon incubation at 28°C for 24 hours in the dark.

### Growth conditions and tissue collection for transcriptome and metabolome profiling

Four day old seedlings of approximately the same radicle length were transferred to a soil plug, that is, a 50 mL falcon tube filled with sterile soil (soil collected from the Clue Field in the Netherlands; 52° 03’ 37.91’’ N and 5° 45’ 7.074’’ E, which was further dried, sieved through 4 mm mesh and sterilized by gamma irradiation) mixed with 5% sterile water (w/v). Seedlings were watered with autoclaved water every second day. On day ten, seedlings together with the soil plug were transferred to 40 cm long cones (Greenhouse Megastore, catalog number CN-SS-DP) filled with 700 mL of 0.5-1.0 mm filter sand (filcom.nl/). Plants were organized in a randomized manner in a greenhouse compartment with a temperature of 28°C during the day (11 hours) and 25°C at night (13 hours), with 70% relative humidity and light intensity of 450μmol/m^2^/s. At days zero, seven and 14, plants were watered with 50 mL of modified half-strength Hoagland solution with 0.05 mM KH_2_PO_4_. On days one, four, ten, 13 and 17, plants were watered with 50 mL deionized sterile water. Two and three weeks after transfer to the cones (corresponding to 28 and 35 day old plants) root material was harvested two hours after the light turned on. Each sorghum plant was gently taken from the cone and the whole root system was cleaned from the sand and soil by washing in water, dried with paper towel and snap frozen in liquid nitrogen (taking approximately 3 minutes per plant).

### RNA-seq library preparation

Root tissue was manually ground and RNA was extracted with RNaesy Plus Mini kit (Qiagen) with application of QIAshredder columns (Qiagen) and on-column Dnase I (Qiagen) treatment. Total RNA obtained was precipitated with 3M NaOAc pH 5.2 (Thermo Scientific) in 100% ethanol and washed with 70% (v/v) ethanol. RNA-seq libraries were synthesized with QuantSeq 3’ mRNA-Seq Library Prep Kit (Lexogen) according to manufacturer protocol. Libraries were sequenced at the UC Davis DNA Technologies Core with Illumina HiSeq 4000 in SR100 mode with four biological replicates and three technical replicates for each RNA sample.

### RNA-seq data processing and differential expression analysis

Quality control of resulting transcriptome sequencing data was accessed with FastQC (http://www.bioinformatics.babraham.ac.uk/projects/fastqc/) before and after read processing. Technical replicates of the libraries were pooled before reads were processed. Barcodes were trimmed from raw reads with fastx-trimmer (http://hannonlab.cshl.edu/fastx_toolkit/index.html) with parameters: -v -f 12 -Q33. Adaptor trimming and quality filtering was performed with reaper from Kraken Suite (Davis *et al*., 2013) with options: -geom no-bc -tabu $tabu -3pa $seqAdapt - noqc -dust-suffix 6/ACTG -dust-suffix-late 6/ACTG -nnn-check 1/1 -qqq-check 35/10 -clean-length 30 -polya 5. Trimmed reads were mapped to the reference genome of *Sorghum bicolor* BTx623 (McCormick *et al*., 2018) using STAR (Dobin *et al*., 2013) with options: -- outFilterMultimapNmax 20 --alignSJoverhangMin 8 --alignIntronMin 20 --alignIntronMax 10000 - -outFilterMismatchNmax 5 --outSAMtype BAM SortedByCoordinate --quantMode TranscriptomeSAM GeneCounts.

Genes whose raw read counts across all samples equaled zero were removed. Counts per million (CPM) were calculated with cpm() function from the edgeR package (Robinson *et al*., 2010) and only genes with a CPM > 1 in a minimum of three samples were used for further analysis. CPM values are found in **Table S1**. Differentially expressed genes (DEGs) were identified with the R/Bioconductor limma package (Ritchie *et al*., 2015). CPM values were normalized with the voom() function with quantile normalization to account for different RNA inputs and library sizes. The linear model for each gene was specified as an interaction of the genotype and the timepoint: log(counts per million) of an individual gene ∼ Genotype*Time. Differentially expressed genes for each term of linear model were selected based on a false discovery rate < 0.05. Lists of differentially expressed genes for each term (genotype, time, genotype-by-time interaction) are found in **Table S2-4**.

### Clustering analysis and ontology enrichment

Genes differentially expressed between genotypes in a time-dependent and time-independent manner were clustered in groups of genes that have similar expression patterns. The 75% most variable genes were selected for clustering. The log_2_ CPM mean across biological replicates were calculated for each gene and expression of each gene was then scaled to the mean expression across all samples. Hierarchical clustering has been performed with pheatmap v.1.0.12 R package with the Euclidean distance measure to quantify similarity. Genes assigned to each cluster are listed in **Table S5**.

Gene Ontology (GO) enrichment analysis was been performed with the GOseq v.1.34.1 R package with correction for gene length bias (Young *et al*., 2010). The odds ratio for each ontology was calculated with the formula: (number of genes in GO category / number of all genes in input) / (number of genes in GO category / number of genes in all clusters). Enriched ontology terms were selected based on a p-value < 0.05 and an odds ratio > 1. Multiple testing correction is not recommended for GO enrichment due to the graph structure of the GO terms (Mi *et al*., 2012). GO categories enriched in each cluster are listed in **Table S6**.

### Metabolomic analysis from 28 and 35 day old plants

Root tissue was manually ground and 70 mg of pulverized tissue was used for the extraction with 0.7 mL 80% methanol (v/v) containing 6.10^−3^ mg/ml Ribitol (Sigma Aldrich, St. Louis, Missouri, United States) as internal standard. The mixture was shaken using a tissuelyzer (Tissuelyzer II, Qiagen) for 5 minutes at 30 hz and subsequently sonicated for 5 minutes. Samples were centrifuged at 7600 rcf in 10 °C for 5 minutes, and the supernatant was collected and filtered through a 0.22 µm pore filter. Metabolite analysis by LC-MS was carried out as described in (Melkonian *et al*., 2020) with slight modifications. Five µL of root extracts were injected on a Nexera UHPLC system (Shimadzu, Den Bosch, The Netherlands) coupled to a high-resolution quadrupole time-of-flight mass spectrometer (Q-TOF; maXis 4G, Bruker Daltonics, Bruynvisweg 16 /18). Compounds were separated on a C18 stationary phase column (1.7 µm particle size, 150 × 2.1 mm; ACQUITY UPLC CSH C18, Waters, EttenLeur, The Netherlands) preceded by a guard column (1.7 µm particle size, 5 × 2.1 mm; ACQUITY UPLC CSH C18, Waters), a flow rate of 0.3 mL min^-1^, and column temperature of 30°C. Gradients of eluent A (0.1% v/v acetic acid in water) and eluent B (0.1% v/v acetic acid in 100% acetonitrile) were as follows: 0-1 min (5% B); 1-15 min (linear increase to 100% B); 15-18 min (100% B). Compound ionization was carried out by ESI operating in the negative mode using N_2_ as the ionization gas with the following settings: capillary voltage 3,500 V; end plate offset 500 V; nebulizer gas pressure (N_2_) 1 bar; dry gas (N_2_) 8 L min^-1^; dry temperature 200°C. Settings for MS analysis were: funnel radio frequency (RF) 200 Vpp (voltage point to point); multipole RF 200 Vpp; collision cell RF 200 Vpp; transfer time 40 µs; prepulse storage 5 µs. Internal mass calibration was performed automatically during every measurement by loop injection of 20 µL of a 2 mM sodium acetate solution in 1:1 v/v ultrapure water-isopropanol. Data acquisition was done using the Bruker Daltonics software suites Compass 2.7. Peak finding, peak integration and retention time correction were performed using the xcms R package version 1.38.0 (Smith *et al*., 2006) after conversion of the original RAW files to an open-source format ‘mzXML’ with the MSconvert tool from ProteoWizard version 3.0.5163 (Adusumilli and Mallick, 2017). The peak picking was performed using the “centwave” method with the following parameters: Signal/Noise threshold = 10, peakwidth = c (2, 24), mzdiff = 0.001, prefilter = c (3, 100). The “obiwarp” method was used for retention time adjustment. Finally, feature correspondence was achieved with the “density” method using the following optimized parameters: bw = 5.0 and mzwid = 0.017. For pathway analysis .mzXML were imported into the xcms online web environment (Forsberg *et al*., 2018) and analyzed using the same parameters as above described. The sample BioSource was set to *Arabidopsis thaliana* var. Columbia for pathway annotation.

### Root system architecture quantification from 28 and 35 day old plants

Sorghum plants were gently taken from the pots and roots were cleaned from the sand and soil by washing in water. Crown roots were separated from the seminal roots and fresh weight was scored for them separately. Roots were then placed in a water tray and scanned at 800dpi resolution with Epson Perfection V700 scanner. Next, roots were dried with a paper towel, placed in paper bags, dried for 48 hours at 65°C and weighed to determine their dry weight. Root scans were analyzed with the DIRT (Digital Imaging of Root Traits) software v1.1 (Das *et al*., 2015). We used DIRT’s area trait that counts the number of pixels representing the root in the image (Bucksch *et al*., 2014) as a measurement for total root network area (to simplify we refer to it as total root length). Total network length (to simplify we refer to it as total root length) was computed by modifying the original code. As such, we count the number of pixels belonging to the medial axis of the network area after removing spurious medial branches whose medial circle radius at the tip is two pixel or smaller (Bucksch, 2014). Mean root network diameter was then calculated as the ratio of network area over network length. Values for each trait were transformed with natural logarithm-transformation. A two-way ANOVA was used to determine the significance of the differences between genotype and genotype-by-time interactions with the following model: lm(trait ∼ genotype * time) followed by a pairwise comparison with the formula: emmeans(model, specs = genotype * time, adjust = “sidak”), with emmeans v.1.5.2-1 R package.

### Root system architecture quantification from seedlings

Germinated seeds of approximately the same radicle length were transferred to 25 cm long (big pouches for SQR and SRN39 comparisons) or 18 cm long (medium pouches for SQR and all *lgs1* genotypes) germination pouches (PhytoAb Inc., catalog number: CYG-38LG/CYG-98LB) filled with 17 mL or 50 mL autoclaved water, for medium and big pouches, respectively. Pouches were placed in the greenhouse with a maintained temperature of 26°C and daylight of approximately 15 hours. Pouches were scanned daily from the second to the seventh day after germination, approximately eleven hours after the start of the light period. The root system architecture of SQR and SRN39 was quantified at each time point imaged, while for the other *lgs1* genotypes (Framida, Birhan, Gobiye), root system architecture was only quantified in seven day old seedlings. Main root angle (MRA), main root length (MRL), lateral root number (#LR) and the length of individual lateral roots (LRs) were quantified by manual tracing with ImageJ. Lateral root density (LRD) was calculated as the ratio of lateral root number to main root length; lateral root length (LRL) as a sum of the lengths of individual lateral roots, average lateral root length (aLRL) as the ratio of lateral root length to the number of lateral roots; total root size (TRS) as a sum of main and lateral root lengths. Prior to statistical analysis, data was transformed (square root-transformation for the number of lateral roots, natural logarithm-transformation for all the other traits, while the main root angle values were not transformed). The differences between genotypes at each individual timepoint was assessed with an ANOVA including individual plant as a random factor with a formula: lmer(trait ∼ Genotime + (1|Rep), data=x, REML=TRUE), where “Genotime” indicates the combination of genotype and a timepoint (lme4 v.1.1-21 R package) followed with custom contrast comparison for two genotypes per each timepoint with a formula: emmeans(model, specs = ∼ Genotime, adjust = “sidak”), where “Genotime” indicates the combination of genotype and a timepoint (emmeans v.1.5.2-1R package). Seven to ten seedlings were used per genotype as biological replicates.

### Exudate collection and strigolactone quantification

Seeds were surface sterilized by agitating in a solution containing 4% (v/v) sodium hypochlorite and 0.2% Tween-20 (v/v) for 30 minutes followed by 3 alternated washes with 70% ethanol (v/v) and sterile water. The disinfected seeds were thoroughly rinsed five times with sterile water and germinated on wet Whatman paper (grade 1) upon incubation at 28°C for 48 hours in the dark. Germinated seeds with approximately same radicle size were transferred to a 50 mL tube filled with washed river sand and perforated at the bottom to allow draining. The falcon tubes were covered with aluminum foil to prevent the roots from being exposed to light. The plants were grown for 14 days with a temperature of 28°C during the day (11 hours) and 25°C at night (13 hours), with 70% relative humidity and light intensity of 450μmol/m^2^/s. Seedlings were watered with modified Hoagland solution with 0.05 mM KH_2_PO_4_. Root exudate was collected from six plants for each of the sorghum genotypes. Each tube was flushed with 5% ethanol (v/v) in water to collect 35 mL of the flow-through. Each exudate sample was purified using solid phase extraction (SPE) with C18 Discovery® cartridges (bed wt. 500 mg, volume 6 mL, Merck). Cartridges were activated using 5 mL acetone and washed with 5 mL distilled water. Twenty milliliters of sample was loaded on the cartridge, that was further washed with 5 mL distilled water. Finally, compounds were eluted using 3 mL acetone. The acetone was evaporated using a SpeedVac (Scanvac, Labgene) and residual water was removed using freeze drying (Heto Powerdry LL1500, Thermo). The sample was reconstituted in 150 µL 25% (v/v) acetonitrile and filtered using a micropore filter (0.22 µm, 0.75 ml, Thermo scientific) prior to ultra-high performance liquid chromatography – tandem mass spectrometer (UHPLC–MS/MS) analysis as described in (Flokova *et al*., 2020). Values for each compound abundances were transformed with natural logarithm-transformation. One-way ANOVA was used to determine the significance of the differences between genotypes with the following model: lm(trait ∼ genotype) followed by a Tukey post-hoc test (adjusted p-value<0.05) with agricolae v.1.3-1 and multcompView v.0.1-8 R packages.

### Growth analysis

Germinated seeds with approximately the same radicle length were transferred to pots containing a custom potting mix (one part coarse sand, one part compost, one part peat and 4.5 lbs/yd dolomite lime) and placed in the greenhouse with a maintained temperature of 26°C and day light of approximately 15 hours. For each individual plant, every second day, starting from day 4, the number of leaves, and plant height (measured from the soil to the bend of the oldest leaf) was counted. The time to reach vegetative growth stages as defined in (Vanderlip and Reeves, 1972) was scored. At the boot stage of each individual plant, when the boot length was approximately 20 cm, the shoot tissue was harvested and separated into leaves and stalk, roots were excavated from the soil and washed, and the fresh weight of each tissue was quantified. The tissue was then dried at 60°C for 7 days and its dry weight measured. Ten plants with each plant as a biological replicate were tested for each genotype.

### Statistics for growth analyses

The time required to reach individual growth stages was statistically assessed with a Wilcoxon test. Differences in biomass between genotypes were assessed using a Welch t-test. The increase in height of each plant was calculated as the slope of a linear curve fitted with a sliding window of 30 cm. The differences between genotypes in height or in the increase in height was assessed with a Welch t-test. The differences between genotypes in height per individual emerged leaf were assessed with an ANOVA including individual plant as a random factor with formula: lmer(height ∼ Genoleaf + (1|Pot), data=x, REML=TRUE), where ‘Genoleaf’ indicates genotype and leaf number combination (lme4 v.1.1-21 R package) followed with a custom contrast comparison for two genotypes for individual leaf number with formula: emmeans(model, specs = ∼ Genoleaf, adjust = “sidak”), where ‘Genoleaf’ indicates genotype and leaf number combination (emmeans v.1.5.2-1 R package).

## Results

### SRN39 has altered expression of genes involved in metabolism and stress responses relative to Shanqui Red

Shanqui Red (SQR) and SRN39 are Sorghum varieties commonly used as Striga susceptible and resistant models, respectively. The resistance to Striga in SRN39 is associated with a mutation in the *LGS1* locus resulting in altered strigolactone composition in the root exudate (Gobena *et al*., 2017). Sequencing of the root transcriptome of Sorghum varieties with the *lgs1* allele (SRN39) revealed increased strigolactone biosynthesis gene expression relative to SQR (Bellis *et al*., 2020). These expression data suggest that next to the differences in the SRN39 root exudate profile, strigolactone levels may also be altered in SRN39 plant tissues. Strigolactones play a role in several aspects of plant growth and development (Brewer *et al*., 2013) and interface with other hormone signaling pathways (Cheng *et al*., 2013). This likely difference in SRN39 whole plant strigolactone composition as well as extensive genetic divergence between SRN39 and SQR suggest that additional differences in SRN39 growth and development should be characterized and considered in the interpretation of experiments using SRN39 and SQR as controls.

To gain insight into biological processes that may differ between these genotypes, we profiled the transcriptomes of 28 and 35 day old plants, the approximate age used for sorghum strigolactone profiling (Gobena *et al*., 2017; Mohemed *et al*., 2018). As anticipated, given the extensive genotypic differences between these varieties (Gobena *et al*., 2017), multidimensional scaling revealed a clear separation of transcriptional landscapes between the two genotypes, as well as an effect of plant age (**Fig. 1A**). Plant age influences the transcriptome of SRN39 roots to a greater extent than in SQR (**Fig. 1A**). We confirmed higher expression of several sorghum strigolactone biosynthetic genes in SRN39 compared to SQR **(Fig. S1)**, as previously observed by Bellis et al. (2020). Given the clear role of sorghum genotype and developmental age (time) in transcriptome variation, we used an ANOVA to identify genes whose expression differs depending on these factors (genotype, time and a genotype-by-time interaction; **Table S2-4**). First, we identified genes whose expression was affected by the sorghum genotype, by time and in a time by genotype interaction (FDR threshold 0.05; **Table S2-4**). To further characterize these differentially expressed genes and their functions, we performed hierarchical clustering of genes affected by genotype itself or by a genotype-by-time interaction, resulting in identification of six co-expressed gene groups.

**Figure 1.**
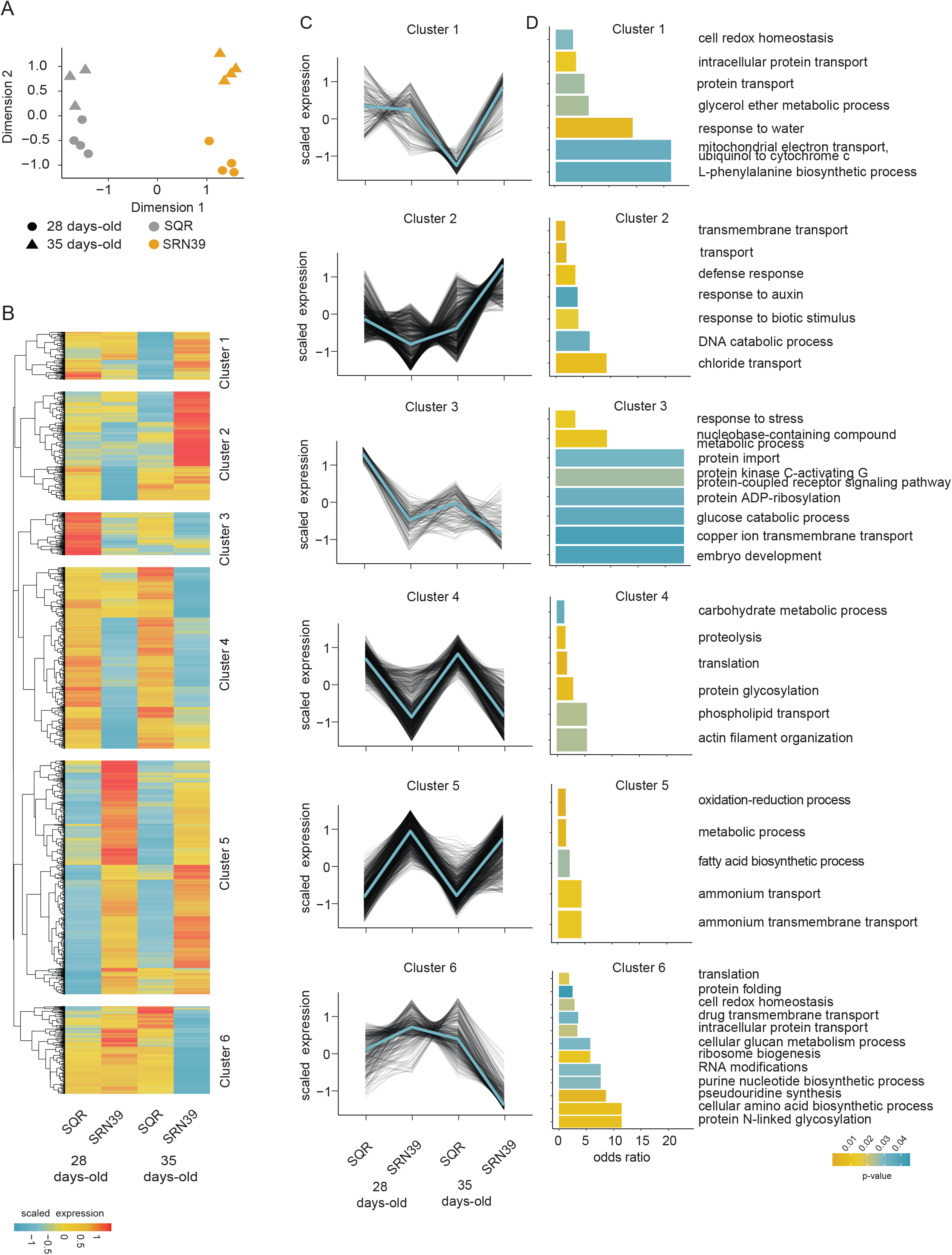
Transcriptome profiles of roots of Shanqui Red (SQR) and SRN39. (A) Multidimensional scaling (MDS) plot of samples based on their transcript abundances (log_2_ Counts Per Million (CPM)). Gray=SQR, orange=SRN39, circle=28 day old, triangle=35 day old. (B) Six clusters of genes affected by a genotype in a time-dependent and independent-manner and (C) corresponding expression patterns. Values presented are log_2_(CPM) scaled to the mean expression across all samples. (D) Biological processes enriched in each cluster. The color of the bar indicates the p-value of the enrichment (at p-value < 0.05 threshold). Root transcriptomes of four biological replicates were sequenced per genotype at each timepoint.

Genes in clusters four and five had similar expression at both time points in each genotype, but different respective expression between genotypes. In cluster four, genes were expressed at lower levels in SRN39 relative to SQR while genes in cluster five, had a higher magnitude of expression in SRN39 relative to SQR. Gene ontology enrichment (p-value < 0.05 and odds ratio > 1) suggests that these genes are associated with metabolism. For instance, in cluster four, where expression is higher in SQR than SRN39, genes are associated with “carbohydrate metabolic process” and “phospholipid transport”. In cluster five, genes are associated with the ontologies “metabolic process”, “fatty acid biosynthetic process” and “ammonium transport” **(Fig. 1C-D)**. In contrast, expression of genes in cluster three decreased over time in both genotypes. Processes enriched within cluster three included “response to stress”, “glucose catabolic process” and “copper ion transmembrane transport” **(Fig. 1C-D)**.

Genes with a more complex genotype-by-time interaction effect were found in clusters one, two and six. Genes from cluster one, associated with “response to water” and “L-phenylalanine biosynthesis process” were more highly expressed in SRN39 at the later timepoint only (**Fig. 1C-D**). Genes with lower expression in SRN39 in 28 day old plants but higher expression in 35 day old plants (cluster two) were enriched in “defense response” and “response to auxin”. Upon further inspection of the genes associated with these clusters we found several genes related to auxin synthesis and transport: *Sobic*.*007G191400, Sobic*.*002G259100*, and *Sobic*.*004G156300* with homology to rice *OsSAUR50 (SMALL AUXIN UP RNA 50), OsSAUR72* and *OsSAUR72-like*, respectively (**Table S5**). Strigolactones regulate polar auxin transport (Cheng *et al*., 2013; Ruyter-Spira *et al*., 2011) and our results suggest this regulation might be dependent on plant age. Genes involved in abscisic acid signaling, *Sobic*.*003G354000* and *Sobic*.*002G172000*, both with homology to Arabidopsis *HAI3 (HIGHLY ABA-INDUCED PP2C PROTEIN, At2g29380*, **Table S5)** were found among those influenced by a complex genotype-by-time interaction (**Fig. 1C-D)**. It is possible that altered strigolactone profiles in SRN39 plants could influence abscisic acid (ABA) signaling. Similarly, strigolactone biosynthetic and signaling mutants had been previously found to have decreased sensitivity to ABA (Ha *et al*., 2014). SRN39 is grown predominantly in Sudan, often in non-irrigated fields. Thus, future research should focus on assessment of SNR39 responses to ABA in relation to drought responses, with consideration of plant age in this regulation.

### SRN39 has altered gibberellin and fatty acid biosynthesis relative to Shanqui Red

The perturbations in expression of metabolism-related genes that we observed in SRN39 roots prompted us to profile the root metabolomes of SQR and SRN39 grown in the same set up as for the transcriptome analysis. Similar to the transcriptome landscapes (**Fig. 1A**), the metabolite profiles of 28 and 35 day old root of SQR and SRQ39 were clearly separated and plant age had a more profound effect on metabolites of SRN39 than SQR (**Fig. 2A**). The metabolic features differentially accumulated between genotypes at both plant ages assessed, were enriched with gibberellin biosynthesis (**Fig. 2B-C)**. Several metabolic features predicted to be gibberellin precursors (ent-kaur-16-en-19-oate, ent-7α-hydroxykaur-16-en-19-oate, gibberellin A12-aldehyde and gibberellin A12) had a higher abundance in roots of SRN39 than in SQR (**Fig. 2D-H**).

**Figure 2.**
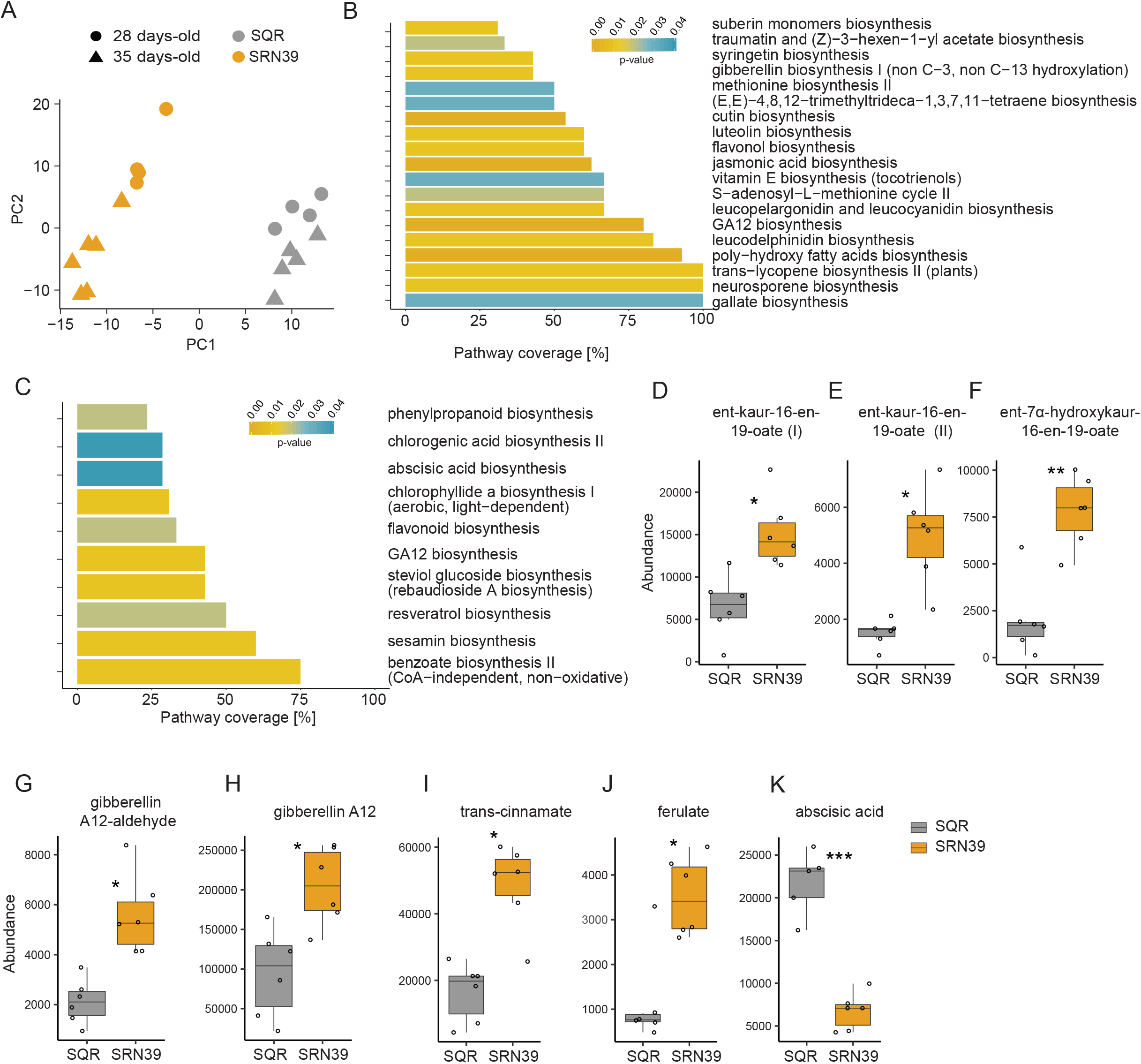
Metabolome profiles of roots of Shanqui Red (SQR) and SRN39. (A) Principal component analysis (PCA) of samples based on their metabolite abundance. Gray=SQR, orange=SRN39, circle=28 days old, triangle=35 days old. Pathway enrichment analysis of metabolites differentially accumulated between genotypes at (B) 28 day old and (C) 35 day old stage. The color of the bar indicates the adjusted p-value of the enrichment (at a < 0.05 threshold), while the x-axis indicates the proportion of the metabolites in each pathway that was found to be differentially accumulated in SRN39 as compared to SQR. Abundance of gibberellin biosynthesis intermediates: metabolic features predicted to be (D, E) ent-kaur-16-en-19-oate, (F) ent-7α-hydroxykaur-16-en-19-oate, (G) gibberellin A12-aldehyde and (H) gibberellin A12; suberin intermediates: (I) trans-cinnamate and (J) ferulate; and (K) abscisic acid; and. Data in D-J present metabolite levels in roots of 28 day old plants, while in K of 35 day old plants. The boxplots denote data spanning from the 25th to the 75th percentile and are centered to the data median. Dots represent individual values. Asterisks denote a significant adjusted p-value for differences between genotypes by Student’s t-test. *** p-value < 0.001, ** p-value < 0.01, * p-value < 0.05, (n=6).

While we observed enrichment of genes associated with fatty acid biosynthesis to be expressed higher in roots of SRN39 than SQR in both 28 and 35 day old plants, metabolites involved in poly-hydroxy fatty acid biosynthesis and suberin monomer biosynthesis were perturbed only in 28 day old roots. We observed higher levels of metabolic features predicted to be ferulate and trans-cinnamate in roots of SRN39 as compared to SQR (**Fig. 2I-J)**. Ferulate and other phenylpropanoids are components of aliphatic suberin. In the roots of 35 day old plants a lower abundance of abscisic acid (**Fig. 2K)** complements observations of differential transcript accumulation of genes associated with ABA signaling. Collectively, our transcriptome and metabolome analysis demonstrate differences in hormonal balance between SQR and SRN39.

Differences in levels of gibberellin precursors and identification of complex gene expression interactions between 28 and 35 day old SQR and SRN39 plants led to the hypothesis that complex differences in their plant growth may also be observed. Observation of the plants we sampled for transcriptome analysis suggest that such differences may occur. While most of the 28 day old SQR and SR39 had five leaves, almost half of 35 day-old SRN39 plant progressed to the six leaf stage, and two-thirds of SQR plants remained in the five leaf stage (**Fig. S2B)**. The SRN39 plants were shorter than SQR at both plant ages assessed (**Fig. S2A**).

### SRN39 root system architecture is distinct from Shanqui Red

Strigolactones were previously described to play a role in root growth (reviewed in (Rasmussen *et al*., 2013)). Given the distinct strigolactone root exudate profiles of SQR and SRN39 and the extensive genotype-by-time interactions in root gene expression between these genotypes, we asked whether this results in different root system architectures between these genotypes. We compared the total root system length and area, mean root network diameter and dry weight of 28 and 35 day old plants grown in the same experimental set up as used for transcriptome and metabolite profiling. Since strigolactones also play a role in the growth of shoot-borne roots (Kohlen *et al*., 2012; Rasmussen *et al*., 2012), we also quantified the above-mentioned traits separately for crown (shoot-borne roots) and seminal roots (roots of embryonic origin). The mean root network diameter of 28 day old SRN39 plants was smaller than that of SQR plants of the same age, whereas in the case of 35 day old plants, SQR displayed a higher mean root network diameter **(Fig. S2F)**. No significant differences were observed in the length and area of seminal roots, crown roots and total root system **(Fig. S2C-N)**. These subtle time-dependent differences in root system architecture resemble the differences in gene expression observed between SRN39 and SQR occurring between the 28^th^ and 35^th^ day of growth (**Fig. S2B, Fig. 1D-E)**. The dry weight of seminal roots, but not crown roots or total root system size was lower in case of SRN39 than in SQR, independent of plant age **(Fig. S2C, G, K)**

In Arabidopsis, strigolactones are associated with the growth of first order lateral roots (Ruyter-Spira *et al*., 2011). Thus, to further characterize this early stage of SRN39 and SQR root system development, we quantified several root traits in SRN39 and SQR seedlings, from the second to seventh day after germination without nutrient supplementation. The SQR main root length was shorter than SRN39 from the very first days after germination and this difference increased over time **(Fig. 3A)**. The main root of SQR deviated more from the gravity vector than the main root of SRN39 **(Fig. 3B)**. While lateral root density was lower in SRN39 than in SQR from the fifth day after germination onwards **(Fig. 3C)**, lateral root length was greater in SRN39 from the third day after germination **(Fig. 3D)**. Although the increase in lateral root length was stable across multiple experiments, the magnitude of the difference in main root length was more variable **(Fig. 3A, D, Fig. 4A-B)**. Our observations suggest therefore, that SRN39 has a longer and steeper main root, and longer lateral roots, compared to SQR at the seedling stage. Together this contributes to a greater total root system size of SRN39 as compared to SQR (**Fig. 3E)**.

**Figure 3.**
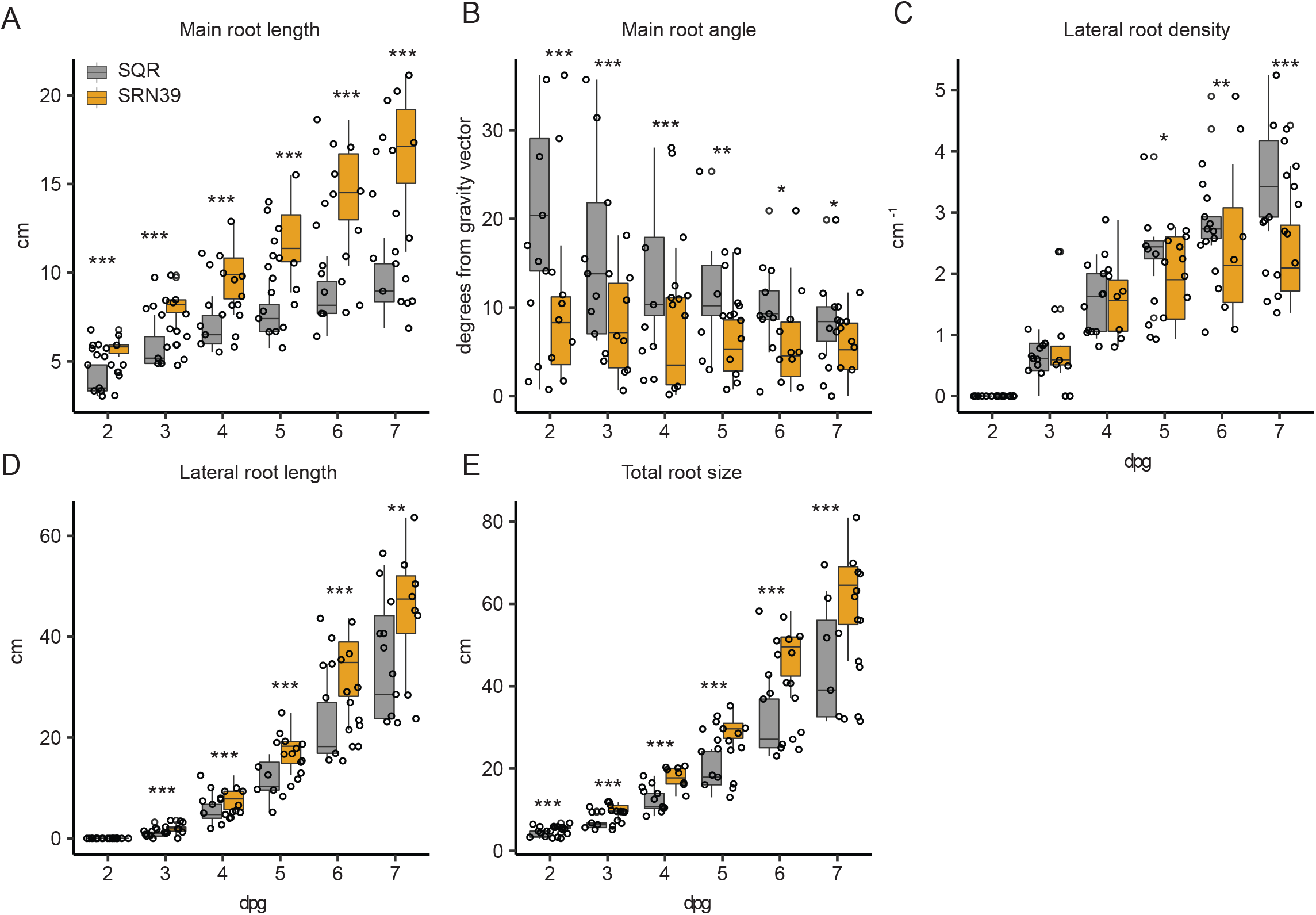
Root system architecture of Shanqui Red (SQR) and SRN39 and sorghum varieties with the *lgs1* mutation. In all cases the x-axis denotes days post germination (dpg). (A-E) SRN39=orange; SQR=gray. (A) Main root length, (B) main root angle, (C) lateral root density, (D) lateral root length, (E) total root size. The boxplots denote data spanning from the 25th to the 75th percentile and are centered to the data median. Dots represent individual values. Asterisks denote a significant p-value for each timepoint between genotypes by the least square method. *** p-value < 0.001, ** p-value < 0.01, * p-value < 0.05, (n=10).

**Figure 4.**
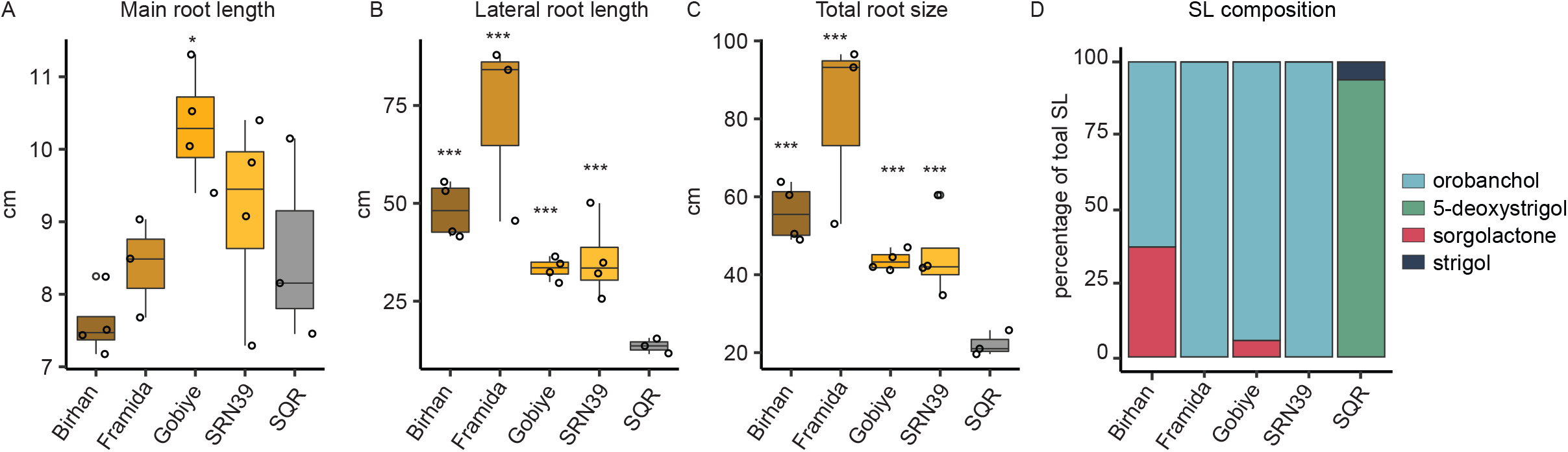
Root system architecture and root exudate strigolactone composition of Shanqui Red (SQR) and sorghum varieties with the *lgs1* mutation (Birham, Framida, Gobiye and SRN39). (A) Main root length, (B) lateral root length, (C) total root size of seven day old seedlings. The boxplots denote data spanning from the 25th to the 75th percentile and are centered to the data median. Dots represent individual values. Asterisks denote a significant p-value for each timepoint between genotypes by the least square method. *** p-value < 0.001, ** p-value < 0.01, * p-value < 0.05, (n=4). (D) Relative abundance (percentage of total strigolactones measured) of four strigolactones in root exudates of 14 day old plants. (n=6). The abundance of individual strigolactones is presented in Figure S5.

### Sorghum genotypes producing orobanchol have increased lateral root growth

To verify whether the observed increase in root system size of SRN39 is determined by the *lgs1* mutation, or by other genetic differences between SQR and SRN39, we evaluated the root system architecture of three other sorghum varieties carrying a mutated *LGS1* variant (Birhan, Framida and Gobiye). The increased main root length was only found in Gobiye **(Fig. 4A)**. All *lgs1* mutant varieties had an increase in average lateral root length **(Fig. S4B)**, total lateral root length (**Fig. 4B)** and total root system length **(Fig. 4C)** relative to SQR. We also confirmed that, similar to SRN39, the *lgs1* mutation in Birhan, Framida and Gobiye varieties resulted in exuding orobanchol as a main strigolactone (**Fig. 4D, Fig. S5**). We suggest that impaired LGS1 function may be associated with the promotion of lateral root growth.

### SRN39 is developmentally delayed, but accumulates more biomass

To evaluate whether the observed differences between SRN39 and SQR root transcriptomes and system architecture are concomitant with distinct growth and development of the above-ground tissue, we quantified SRN39 and SQR vegetative growth in pots under standard greenhouse conditions over time. SRN39 plants were shorter than SQR, with the largest differences observed within the first 30 days of growth, after which the differences in plant height diminished, to increase again around day 70 (**Fig. 5A**). Most of the SRN39 plants did not reach the height of SQR at their respective boot stage (**Fig. 5A)**. To determine whether the differences in height between SQR and SRN39 observed at the boot stage are caused by a decrease in growth specifically within early developmental stages we calculated the rate of height increase given individual plant height. SQR and SRN39 had a distinct growth rate for plants for heights up to 40 cm and for heights greater than 100 cm indicating that there are two phases of growth slow-down in SRN39 (**Fig. 5B**).

**Figure 5.**
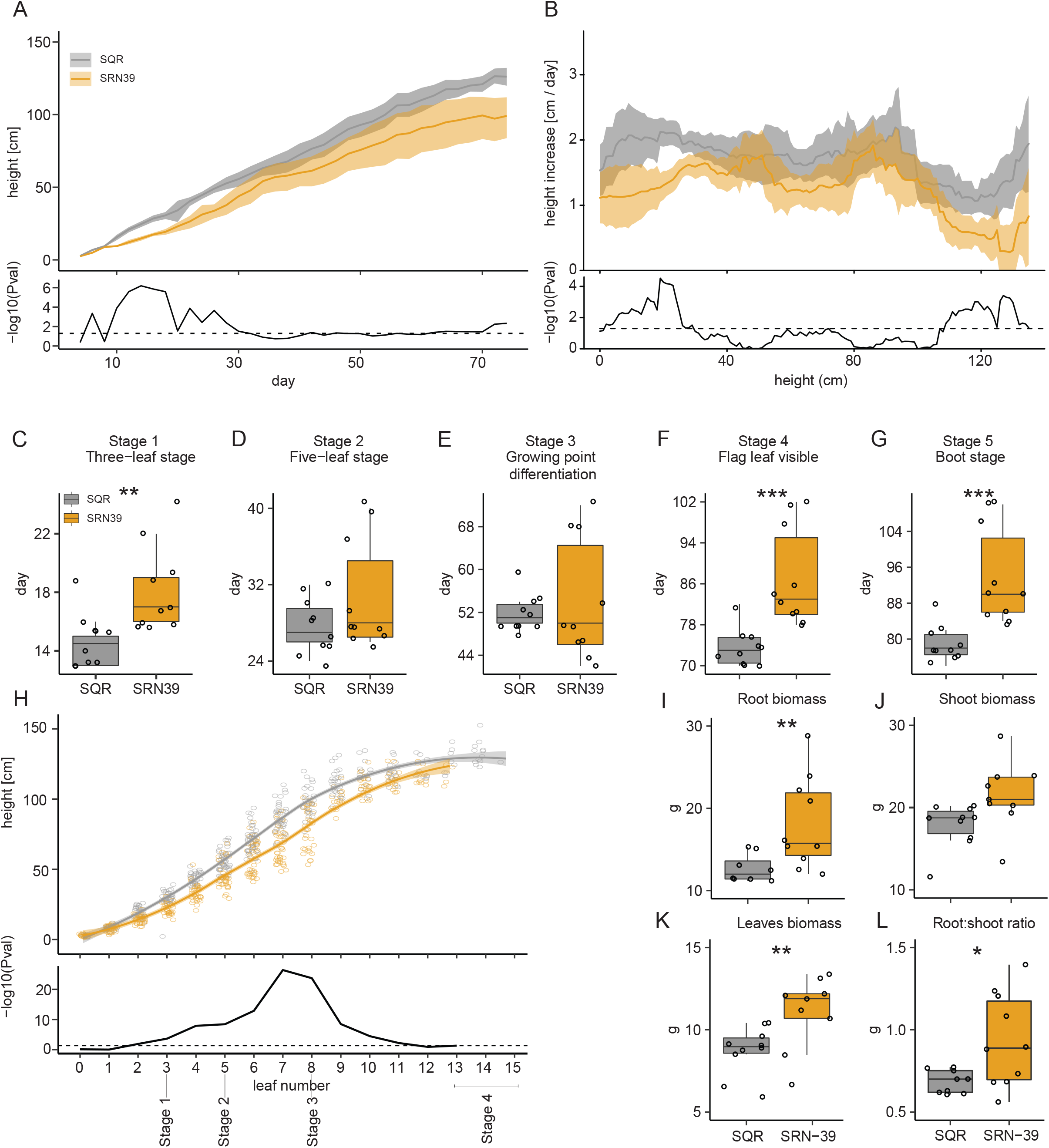
Vegetative growth of SRN39 and Shanqui Red (SQR). (A) Plant height from day four to 74. (B) Increase in plant height per plant height calculated with a sliding window of 30 cm. Data presented include the mean (solid line) and the 95 % confidence interval (shaded). The bottom panel denotes the -log_10_(p-value) at each timepoint (Welch t-test). The dashed line indicates a p-value of 0.05. Time required to reach (C) stage 1 (three-leaf stage) (D) stage 2 (five-leaf stage), (E) stage 3 (growing point differentiation), (F) stage 4 (flag leaf visible), (G) stage 5 (boot stage). An asterisk denotes the p-value from a Wilcoxon test. (H) Plant height at individual leaf stages. The mean is represented by a solid line, while the 95 % confidence interval is shaded. The bottom panel denotes -log_10_(p-value) per leaf stage as a result of pairwise genotype comparisons using the least square method. (I) Dry weight of root, (J) stalk, (K) leaves, and (L) ratio of root to shoot dry weight. Boxplots denote the span from 25th to the 75th percentile and are centered to the data median. Dots represent individual values. Asterisk denotes p-value from Welch t-test. *** p-value < 0.001, ** p-value < 0.01, * p-value < 0.05, (n=10).

We next compared the time required for each genotype to reach specific stages of sorghum vegetative growth (Vanderlip and Reeves, 1972). Differences between SRN39 and SQR were observed at stage 1 (three-leaf stage), stage 4 (flag leaf visible) and stage 5 (boot stage). No differences were observed between SQR and SRN39 in the time required to reach stage 2 (five-leaf stage) and stage 3 (growing point differentiation) (**Fig. 5C-G**).

To verify whether the growth slow-down and developmental delay are coupled in SRN39, we compared the height of SRN39 and SQR relative to the emergence of each leaf. The differences in plant height between SRN39 and SQR were most prominent in the five-leaf stage (stage 2) and stage 3 (**Fig. 5H)**. Given that there is no difference in the time to reach stage 2 and stage 3 between SQR and SRN39, a developmental delay cannot explain these final differences in plant height (**Fig. 5D-E, H)**. Over the entire vegetative growth period monitored, most SRN39 plants developed a maximum of 13 leaves, while most SQR developed a maximum of 15 leaves (**Fig. 5H)**. These collective data suggest that SRN39 is developmentally delayed at stage 1 only, while the later differences in growth between SRN39 and SQR are not due to a developmental delay.

We further assessed the biomass of the below- and above-ground (leaves and stalk) tissue in SRN39 and SQR. Consistent with the observed increase in RSA of SRN39 compared to SQR, the dry weight of the SRN39 root system was higher than that of SQR (**Fig. 5I)**. No differences were observed in the dry weight of the shoot tissue (leaves and stalk together) (**Fig. 1J)**, while the biomass of leaves was higher in SRN39 compared to SQR (**Fig. 5K)**. The root biomass to shoot biomass ratio was also higher in SRN39 plants compared to SQR (**Fig. 5L)**.

To summarize, while differences in early and late stages of vegetative growth (height) are observed between SRN39 and SQR, these are due to a developmental delay only at Stage 1. A developmental delay is also observed at stages 4 and 5 in SRN39 relative to SQR (**Fig. 5F, G)**. Despite being shorter and having less leaves (**Fig. 5H)**, SRN39 allocates more biomass to its roots and leaves (**Fig. 5I, K)**.

## Discussion

Shanqui Red (SQR) is a sweet sorghum variety originating from China, while SRN39 is a released Sudanese variety. Despite their vast genetic diversity, the differences between these two genotypes have previously been attributed to the polymorphism in the *LGS1* locus leading to the changes in composition of their root exudates (Gobena *et al*., 2017). SQR exudes 5-deoxystrigol as its dominant strigolactone, while SRN39 produces only orobanchol (Gobena *et al*., 2017) (**Fig. 4D)**. 5-deoxystrigol and orobanchol have distinct activity in terms of their stimulation of Striga germination. 5-deoxystrigol is an efficient germination stimulant, while orobanchol only induces poor germination (Gobena *et al*., 2017). Consequently, SRN39 shows resistance to Striga in the field as opposed to the highly susceptible SQR (Gobena *et al*., 2017; Mohamed *et al*., 2003). The predominant role of strigolactone within root exudates, however, is recruitment of arbuscular mycorrhiza fungi, while parasitic plants co-opted strigolactone signalling into their host plant-dependent lifecycle (Aquino *et al*., 2020). Increasing evidence suggests that strigolactones also govern assembly of the rhizosphere microbiome (reviewed in (Aliche *et al*., 2020). Both 5-deoxystrigol and orobanchol have high activity in terms of induction of mycorrhizal hyphal branching (Akiyama *et al*., 2010). Indeed, despite the differences in stereochemistry of the dominant strigolactones in SQR and SRN39, this does not affect their ability to recruit arbuscular mycorrhizal fungi (Gobena *et al*., 2017). On the other hand, increased colonization of bacteria from the Comamondadaceae, Burkholderiaceae and Acidobacteria families has been reported for SRN39 roots (Schlemper *et al*., 2017). Further research is necessary to validate whether the differences in microbial community recruitment are solely the effect of differences in the strigolactone profiles or the consequence of genetic variation other than within the *LGS1* locus.

Next to the emerging evidence for different roles of strigolactone variants in their interaction with biotic agents, the role of strigolactone structural variation in plant development is not well understood. We observed that SRN39 has longer main and lateral roots, and greater overall root length than SQR in young seedlings, while at the end of the vegetative growth SRN39 roots have an increased biomass **(Fig. 2, Fig. 5I)**. The three other sorghum varieties with a *lgs1* mutation, and orobanchol as the dominant exuded strigolactone, also showed longer lateral roots and larger root systems. We cannot distinguish whether this is an effect of the exuded strigolactones on the root or strigolactone composition within the root tissue. Nevertheless, these results suggest that orobanchol has a greater potential to promote lateral root growth than 5-deoxystrigol. Previously, a synthetic strigolactone, GR24, has been shown to repress lateral root formation under phosphate sufficient conditions, while enhancing it under conditions of low phosphate availability (Ruyter-Spira *et al*., 2011). Here we show that, in the absence of nutrients, strigolactones are also involved in the elongation of lateral roots and speculate that different stereoisomers of strigolactones may have distinct potential in shaping the sorghum root system.

The enhanced growth of lateral roots in SRN39 seedlings was reflected in the higher total root biomass of mature plants, as compared to SQR (**Fig. 3 D, Fig. 4I**). The increased investment in root biomass accumulation might impair the growth of the above-ground plant organs (Lynch, 2007). We observed that both the growth and development of SRN39 were affected when compared with SQR. SRN39 was delayed in reaching very early (stage 1, three leaf stage), and late (stage 4 - flag leaf visible, stage 5 - boot stage) stages of vegetative growth and the growth rate (expressed as height) of its above-ground organs was also diminished at these stages (**Fig. 4A-H**). The increased investment in SRN39 root growth was observed at the early seedling stages and at the end of the vegetative growth, thus coinciding with the stages when the growth slow-down of the shoot was the largest and when developmental delay was observed, thus at stage 1 (three leaf stage), stage 4 (flag leaf visible) and stage 5 (boot stage) **(Fig. 2, Fig. 4B,C, F, GI)**, The differences in root system architecture and root biomass were marginal in plants of five to six leaves (28 and 35 day-old plants), thus at the stage where no developmental delay was observed for SRN39 plants and when its growth rate began to increase **(Fig. S1B-N, Fig. 4B, D)**. Together, these data suggest that the increased root growth of SRN39 is linked to both its developmental delay as well as reduced rate of growth above-ground. These differences between SRN39 and SQR could further be attributed to distinct molecular profiles.

The complex relationship between shoot and root growth over time in SRN39 was also reflected in the root transcriptome and metabolic profiles. The effect of plant age on both the transcriptome and metabolome landscape was more profound in SRN39 than SQR (**Fig. 1A, Fig. 2A)**. Among the metabolic pathways perturbed in roots of SRN39 as compared to SQR, gibberellin biosynthesis was observed for 28 and 35 day old plants (**Fig. 2B-C)**. Several gibberellin precursors accumulated to higher levels in SRN39 as compared to SQR, which may be a reason why SRN39 plants are shorter than SQR (**Fig. 2D-H)**. Recently Bellis et al. (2020) created a CRISPR-edited knock-out of *LGS1* in the Macia variety and profiled its shoot transcriptome. Two transcription factors involved in floral initiation, *Sobic*.*010G180200* and *Sobic*.*008G168400* were differentially expressed in the shoot of a CRISPR-edited knock-out of *LGS1* as compared to the wild type, Macia (Bellis *et al*., 2020), which could explain the delay in reaching a boot stage by SRN39 (**Fig. 5G**). Together, these results suggest that the growth slow-down and developmental delay of SRN39 plants may be attributed to decreased biosynthesis of active gibberellins and an increase of its precursors and altered expression of transcription factors controlling floral initiation, respectively.

Genes we identified as differentially expressed between SQR and SRN39 roots were enriched with processes related to metabolism (“carbohydrate metabolic process”, “metabolic process”, “fatty acid biosynthesis”) and responses to biotic stimuli (**Fig. 1C-D)**. Similar processes were enriched among genes differentially expressed in the CRISPR-edited *lgs1* knock-out and its wild type (Bellis *et al*., 2020). Genes related to fatty acid biosynthesis had higher expression in roots of SRN39 than in SQR (**Fig 1C-D**, cluster 5). Consequently, among the metabolic pathways perturbed in SRN39 roots, we found poly-hydroxy fatty acid biosynthesis and suberin monomer biosynthesis (**Fig. 2B-C)**. Suberin, a long chain fatty acid polymer, forms a protective barrier from pathogens (Andersen *et al*., 2015; Holbein *et al*., 2019; Thomas *et al*., 2007) and may serve as an additional barrier from Striga parasitism in SRN39. Indeed, polymorphisms in genes associated with suberin and wax-ester biosynthesis were recently associated with levels of Striga occurrence (Bellis *et al*., 2020), suggesting that, in addition to pre-attachment resistance, SRN39 may provide another layer of protection from Striga parasitism. Genes involved in fatty acid biosynthesis were also enriched among genes differentially regulated in shoot tissue between the CRISPR-edited knock-out of *LGS1* and its wild type (Bellis *et al*., 2020). While SRN39 plants were shorter than SQR, they also accumulated more leaf biomass, which could be due to the increased production of waxes (**Fig. 5A, K**)

SRN39 has not only increased resistance to Striga, but it is also Striga tolerant, meaning it has a reduced number of Striga infections and that successful Striga attachments cause minimal damage (Nasreldin, 2018; Rodenburg *et al*., 2005). Striga tolerance is often measured as the degree of decrease in yield, photosynthesis efficiency or above-ground tissue biomass (Rodenburg *et al*., 2008; Van Ast *et al*., 2000). We observed the dynamic character of the growth slow-down in SRN39, increases in leaf, but not whole shoot biomass and complex gene expression patterns over time that distinguish it from the Striga-sensitive SQR (**Fig. 3C-D, Fig. 5A-B, J-K**). This suggests that accounting for the developmental stage and tissue used to access sorghum biomass might be important for assessing the tolerance of sorghum varieties with previously reported resistance to Striga.

## Supplementary Data

**Supplementary Table 1**. CPM values for root transcriptome profiling.

**Supplementary Table 2**. Genes detected as differentially expressed between genotypes.

**Supplementary Table 3**. Genes detected as differentially expressed between timepoints.

**Supplementary Table 4**. Genes detected as differentially expressed between genotypes in a time-dependent manner.

**Supplementary Table 5**. Genes assigned to expression clusters.

**Supplementary Table 6**. GO terms enriched in each expression cluster.

**Supplementary Table 7**. Intensity values for root metabolite profiling

**Supplementary Table 8**. Abundance of metabolic features of selected enriched categories in roots of 28 day old plants.

**Supplementary Table 9**. Abundance of metabolic features of selected enriched categories in roots of 35 day old plants.

**Dataset S1**. Raw data from the phenotyping experiments

## Acknowledgements

DK, TT, BT, AW, HEV, HB and SMB gratefully acknowledge support from the Bill and Melinda Gates Foundation, Seattle, WA via grant OPP1082853 “RSM Systems Biology for Sorghum”. SMB is partially funded by an HHMI Faculty Scholar grant. AB was in part supported by the USDOE ARPA-E ROOTS Award Number DE-AR0000821 by the NSF CAREER Award No. 1845760 to A.B. Any Opinions, findings, and conclusions or recommendations expressed in this material are those of the author(s) and do not necessarily reflect those of the National Science Foundation. We would like to thank Peggy Lemeaux for her assistance in bulking Shanqui Red seeds, Ludek Tikovsky and Harold Lemereis for assistance in the greenhouse. We acknowledge Rick Helmus and Eva de Rijke (Institute for Biodiversity and Ecosystem Dynamics (IBED), University of Amsterdam) for their help with LC-QTOF analysis.

## Author Contributions

Conceptualization: DK, TT, BT, SMB. Data curation: DK, TT, BT. Formal analysis: DK, BT, AB. Funding acquisition: HB, SMB. Investigation: DK, TT, ZM, HEV, AW, SMB. Methodology: DK, BT, AB. Project administration: HB, SMB. Software: DK, BT, AB. Supervision: DK, BT, HB, SMB. Validation: DK, TT, BT, AW. Visualization: DK. Writing – original draft: DK, BT, AB, HB, SMB.

## Data availability statement

Sorghum sequences were deposited in NCBI GEO under the accession number GSE167101. Raw data from the growth measurements, root system architecture analysis, strigolactone measurements, ANOVA tables and p-values for each statistical test can be found in **Supplementary Dataset 1**. Data analysis scripts are publicly available at https://github.com/DorotaKawa/Sorghum-growth-development.git.

## Abbreviations

ABA: abscisic acid
AM: arbuscular mycorrhiza
LGS1: Low Germination Stimulant 1
SQR: Shanqui Red

---

**Figure S1.**
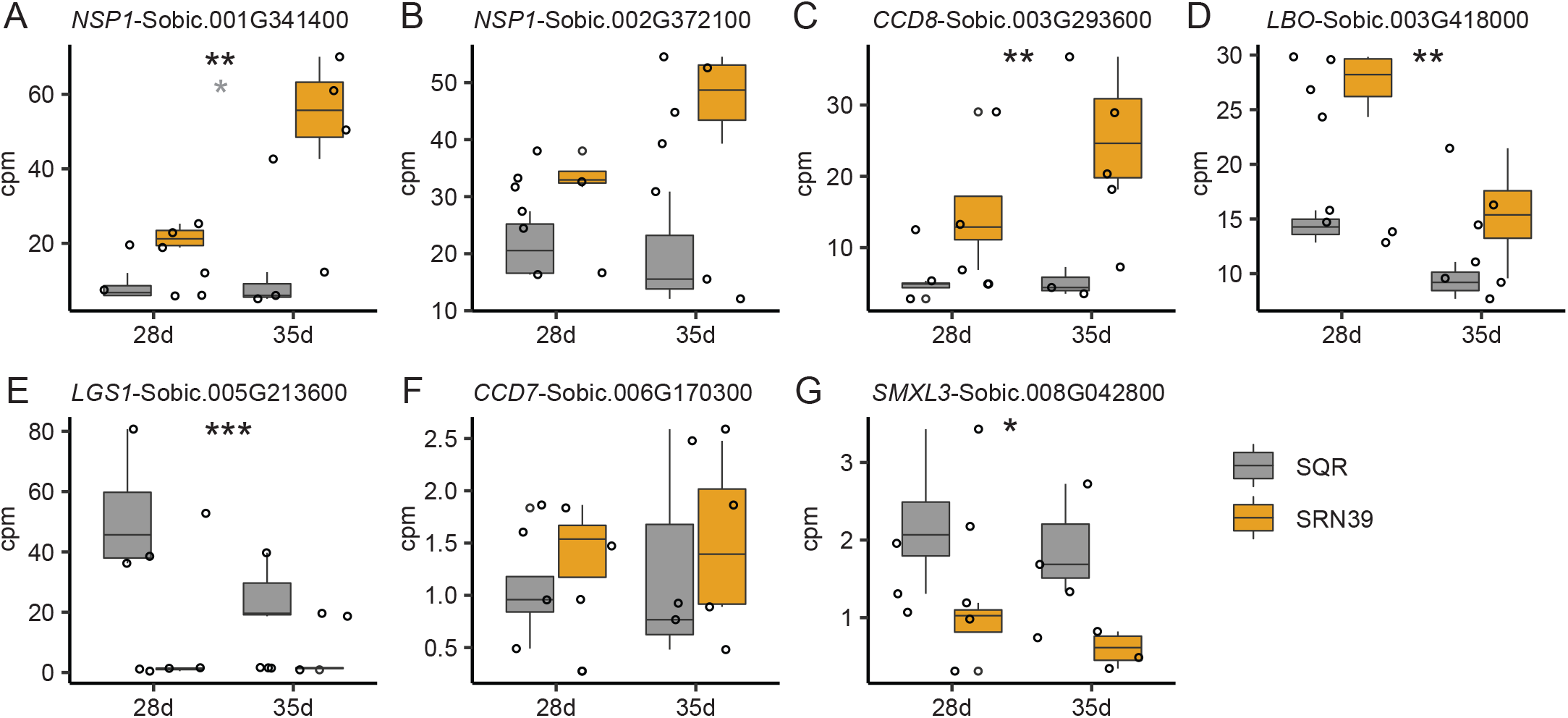
Expression of strigolactone biosynthesis and signaling pathway genes in roots of 28 and 35 day old plants of Shanqui Red (SQR) and SRN39. In all cases the x-axis denotes plant age (28d – 28 day old, 35d – 35 day old plants). (A-B) SRN39=orange; SQR=gray. Transcripts in A, B, D-G has been previously reported as differentially expressed between SQR and SRN39 in (Bellis *et al*., 2020). The boxplots denote data spanning from the 25th to the 75th percentile and are centered to the data median. Dots represent individual values. Black asterisks denote a significant adjusted p-value for term genotype, while grey asterisk indicate significant adjusted p-value for the genotype*time interaction. *** p-value < 0.001, ** p-value < 0.01, * p-value < 0.05, (n=4).

**Figure S2.**
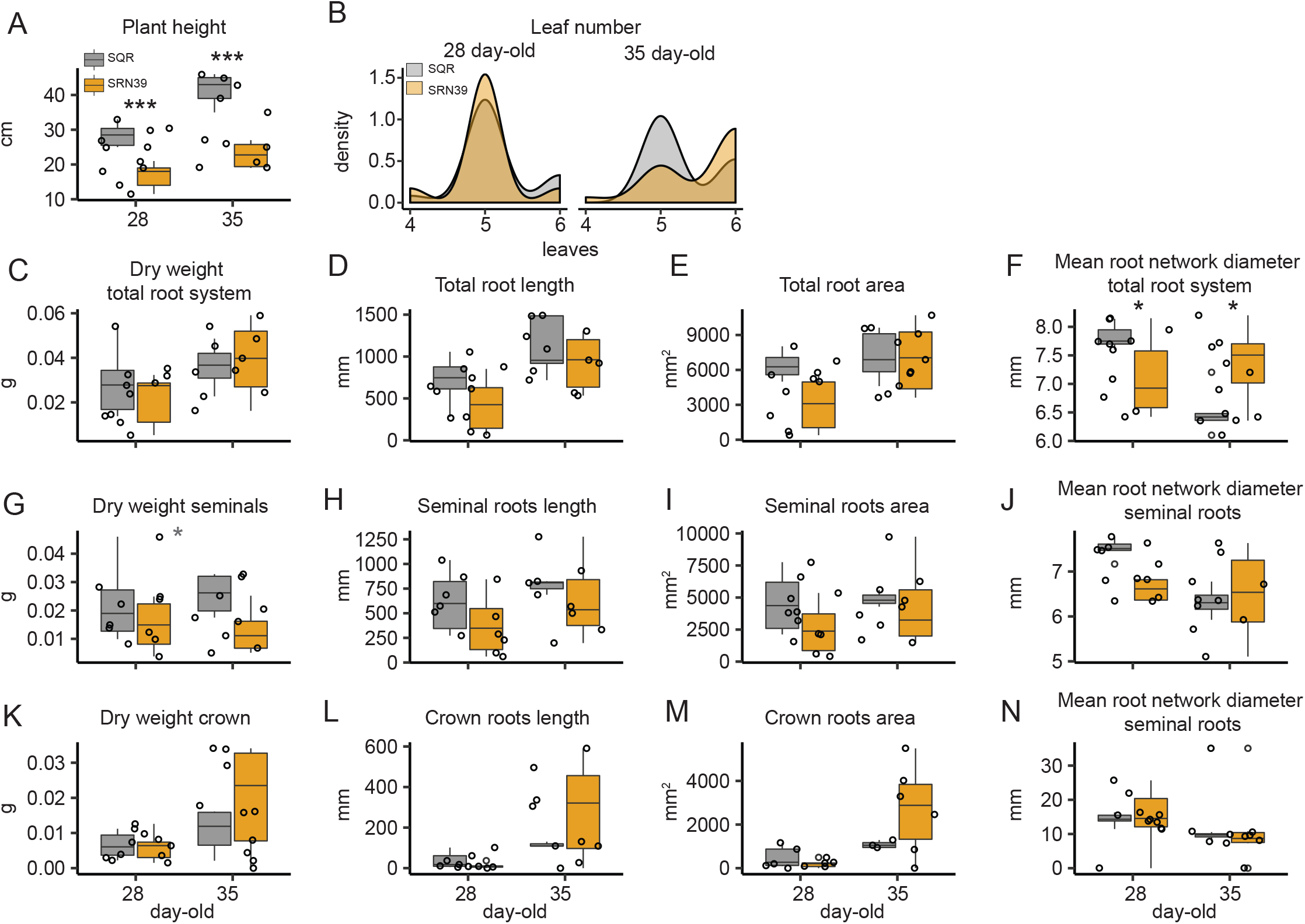
Phenotypic characterization of 28 and 35 day old plants of Shanqui Red (SQR) and SRN39. (A-N) SRN39=orange; SQR=gray (A) Plant height and (B) leaf number, dry weight, root length, root area and mean root network diameter of (C-F) total root system, (G-J) seminal roots and (K-N) crown roots. The boxplots denote data spanning from the 25th to the 75th percentile and are centered to the data median. Dots represent individual values. Black asterisks denote a significant p-value for a post-hoc pairwise comparison between each genotype withing a timepoint by the least square method. Gray asterisks denote a significant p-value for a genotype term according to a two-way ANOVA. *** p-value < 0.001, ** p-value < 0.01, * p-value < 0.05, (n=6 for A, C-N; n= 15-22 for B).

**Figure S3.**
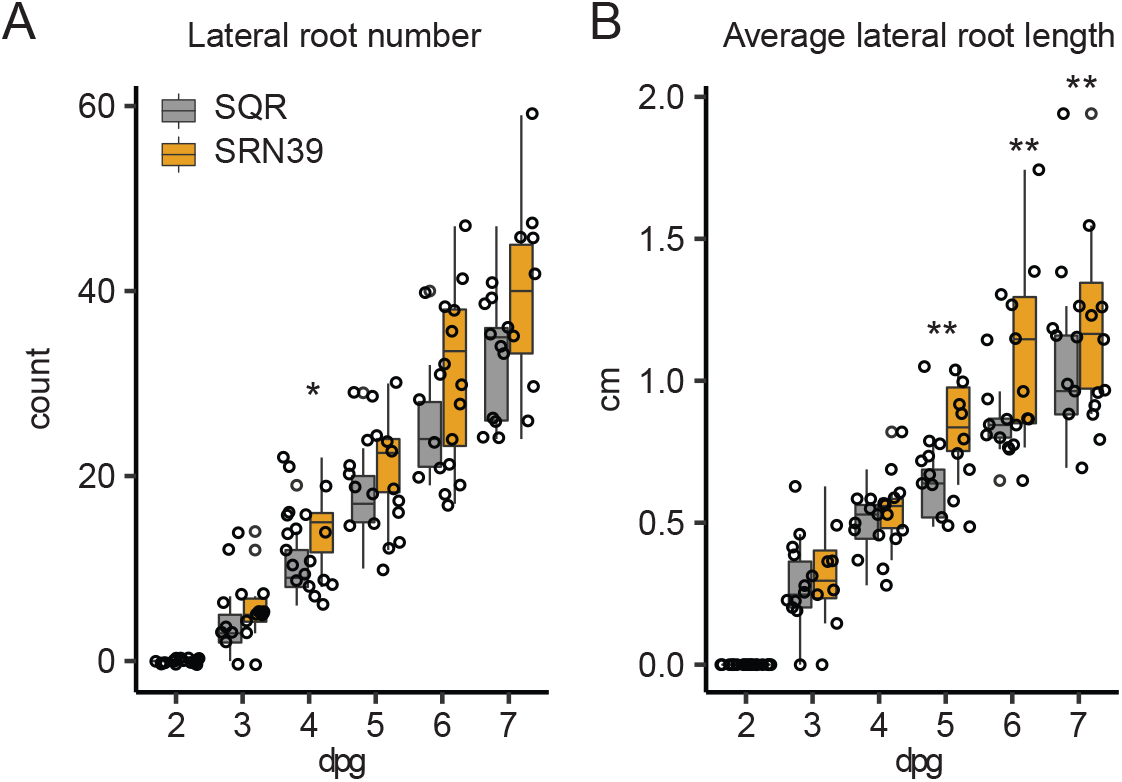
Root system architecture of Shanqui Red (SQR) and SRN39 and sorghum varieties with the *lgs1* mutation. In all cases the x-axis denotes days post germination (dpg). (A-B) SRN39=orange; SQR=gray. (A) Lateral root number, (B) average lateral root length. The boxplots denote data spanning from the 25th to the 75th percentile and are centered to the data median. Dots represent individual values. Asterisks denote a significant p-value for each timepoint between genotypes by the least square method. *** p-value < 0.001, ** p-value < 0.01, * p-value < 0.05, (n=10).

**Figure S4.**
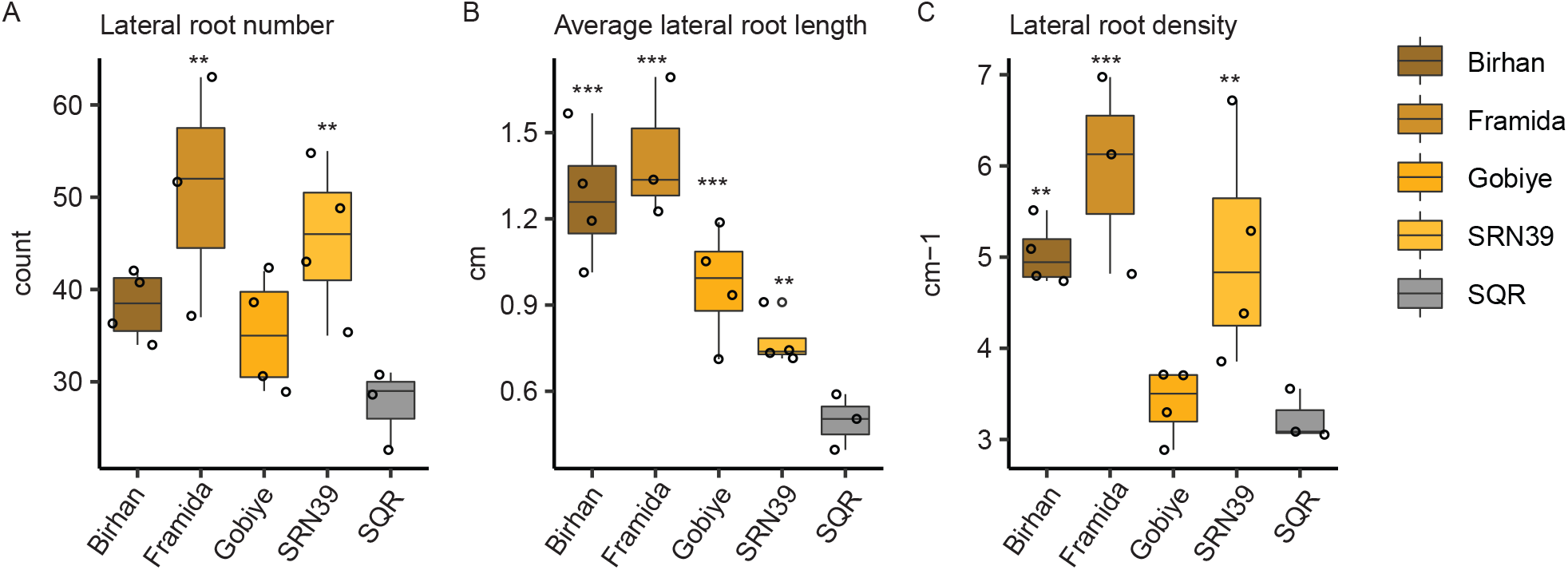
Root system architecture of seven day old seedlings of Shanqui Red (SQR) and sorghum varieties with the *lgs1* mutation (Birham, Framida, Gobiye and SRN39). (A) Lateral root number, (B) average lateral root number, (C) lateral root density. The boxplots denote data spanning from the 25th to the 75th percentile and are centered to the data median. Dots represent individual values. Asterisks denote a significant p-value for each timepoint between genotypes by the least square method. *** p-value < 0.001, ** p-value < 0.01, * p-value < 0.05, (n=4).

**Figure S5.**
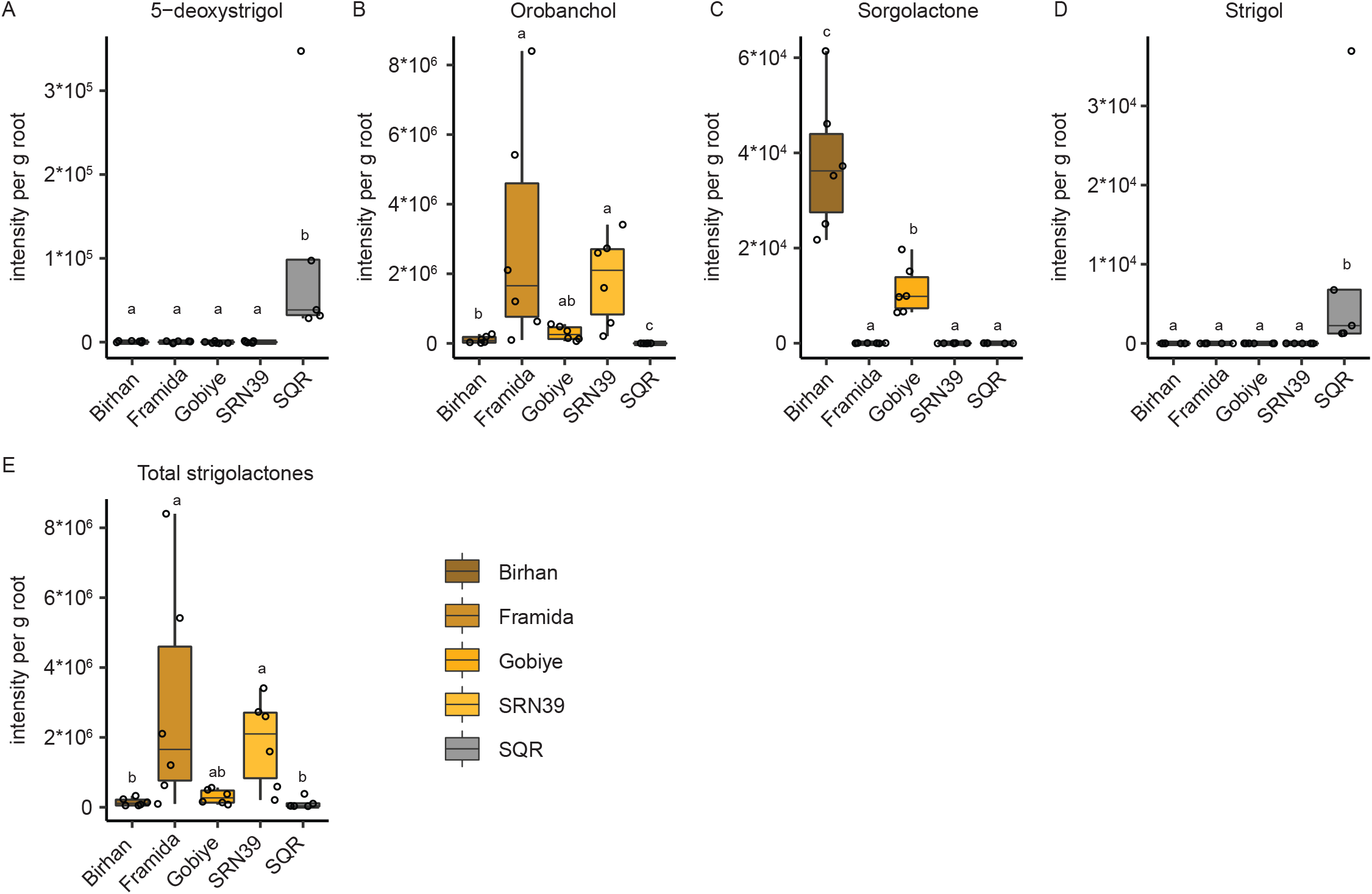
Strigolactone composition in root exudates of Shanqui Red (SQR) and sorghum varieties with the *lgs1* mutation (Birham, Framida, Gobiye and SRN39). Abundance (expressed as intensity per gram of fresh root weight) of (A) 5-deoxystrigol, (B) orobanchol, (C) sorgolactone, (D) strigol and (E) total strigolactones in exudates of 14 day old plants (n=6). The boxplots denote data spanning from the 25th to the 75th percentile and are centered to the data median. Dots represent individual values. Statistical analysis has been performed with one-way ANOVA with Tukey post-hoc test (adjusted p-value<0.05). Different letters indicate significance of difference between tested genotypes.

